# Bottleneck revisited: increased adaptive variation despite reduced overall genetic diversity in a rapidly adapting invader

**DOI:** 10.1101/557868

**Authors:** Daniel Selechnik, Mark F. Richardson, Richard Shine, Jayna DeVore, Simon Ducatez, Lee A. Rollins

## Abstract

Invasive species often exhibit rapid evolution following introduction, despite genetic bottlenecks that may result from small numbers of founders; however, some invasions may not fit this “genetic paradox.” The invasive cane toad (*Rhinella marina*) displays high phenotypic variation across its environmentally heterogeneous, introduced Australian range. Here, we used three genome-wide datasets to characterize population structure and genetic diversity in invasive toads: 1) RNA-Seq data from spleen sampled in the native range (French Guiana), and introduced populations from Hawai’i (the source of the Australian introduction) and Australia; 2) RNA-Seq data from brains sampled more extensively in Hawai’i and Australia; and 3) previously published RADSeq data from transects across Australia. We found that toads form three genetic clusters: (1) native range, (2) Hawai’i and long-established areas near introduction sites in Australia, and (3) more recently established northern Australian sites. In addition to strong divergence between native and invasive populations, we find evidence for a reduction in genetic diversity following introduction. However, we do not see this reduction in loci putatively under selection, suggesting that genetic diversity may have been maintained at ecologically relevant traits, or that mutation rates were high enough to maintain adaptive potential. Nonetheless, cane toads encounter novel environmental challenges in Australia and appear to respond to selection across environmental breaks; the transition between genetic clusters occurs at a point along the invasion transect where temperature rises and rainfall decreases. We identify loci known to be involved in resistance to heat and dehydration that show evidence of selection in Australian toads. Despite well-known predictions regarding genetic drift and spatial sorting during invasion, this study highlights that natural selection occurs rapidly and plays a vital role in shaping the structure of invasive populations.

## Introduction

The genetic paradox of invasion (Allendorf, 2003) describes a phenomenon that challenges widespread evidence of the relationship between genetic diversity and adaptive potential. High genetic diversity within a population is beneficial because it likely underlies phenotypic variation, allowing the population to respond to selection imposed by environmental change (Frankham, 2005; Reed & Frankham, 2003). Furthermore, a greater number of alleles confers an increased frequency of heterozygosity, which is often associated with population fitness (Reed & Frankham, 2003). Small or isolated populations with low genetic diversity have been shown to suffer declines due to inbreeding depression and the associated reduction of individual fitness (Blomqvist, Pauliny, Larsson, & Flodin, 2010; Madsen, Shine, Olsson, & Wittzell, 1999; Westemeier et al., 1998). Conservation efforts to salvage such populations by introducing individuals from allopatric populations (thereby introducing new alleles; “genetic rescue”) have been successful, suggesting that the maintenance of genetic diversity can be crucial for population viability (Madsen et al., 1999; Westemeier et al., 1998).

Despite the fact that invasive populations are thought to undergo genetic bottlenecks due to the translocation of a small number of founders from their native range to an introduced range (Allendorf, 2003; Barrett & Kohn, 1991), invasive species are also characterized by their ability to establish and spread in their introduced ranges. Invasion success is commonly linked to rapid evolution, including adaptation to novel environmental conditions over short timescales (Franks & Munshi-South, 2014; Gao, Li, & Zhan, 2018; Leydet, Grupstra, Coma, Ribes, & Hellberg, 2018; Whitney & Gabler, 2008). Additionally, some invaders exhibit novel phenotypic traits that enhance invasive potential, such as increased growth and dispersal rates (White, Perkins, Heckel, & Searle, 2013; Whitney & Gabler, 2008). There are many examples of evolutionary change during invasion without high levels of genetic diversity (L.A. Rollins, Moles, & Lam, 2013).

Although low genetic diversity may limit the ability of an invasive population to respond to natural selection, rapid evolution can also occur through non-adaptive processes: (1) genetic drift may occur on range edges, reducing genetic diversity across an introduced range (L. A. Rollins, Woolnough, Wilton, Sinclair, & Sherwin, 2009); (2) due to spatial sorting, the invasion front may be inhabited exclusively by the individuals with the highest dispersal rates (even if they are not the fittest) because they have arrived first and can only breed with each other (Shine, Brown, & Phillips, 2011), resulting in a geographic separation of phenotypes (C.M. Hudson, G. P. Brown, & R. Shine, 2016; Shine et al., 2011); (3) admixture or hybridization may occur between individuals from different introductions or sources (Mader, Castro, Bonatto, & de Freitas, 2016).

It has recently been suggested that the genetic paradox of invasion may be rare; some invasive populations do not suffer a reduction in genetic diversity during introduction, and others do not face novel adaptive challenges in their introduced ranges (A. Estoup et al., 2016). Furthermore, some invasive systems demonstrate a ‘spurious’ paradox due to inadequate estimation of genetic diversity (i.e. too few markers), or to maintenance of genetic diversity only at ecologically relevant traits, or to a reduction in genetic diversity resulting from natural selection rather than from genetic bottlenecks (A. Estoup et al., 2016). These ideas can be tested with genome-wide data (Wellband, Pettitt-Wade, Fisk, & Heath, 2018; Willoughby, Harder, Tennessen, Scribner, & Christie, 2018).

The Australian cane toad *(Rhinella marina*) is an invader that has exhibited rapid evolution despite apparently low genetic diversity (L. A. Rollins, Richardson, & Shine, 2015). Cane toads were serially translocated from French Guiana and Guyana to Martinique and Barbados, to Jamaica, to Puerto Rico, to Hawai’i before 101 founders were introduced from the Hawai’ian island of Oahu to Queensland (QLD), Australia in 1935 (Turvey, 2009). Toads have since spread westward through the Northern Territory (NT) into Western Australia (WA; Fig 1). Prior to reaching Australia, toads were exposed to tropical environments with relatively high mean annual rainfall (3200 mm in French Guiana, 1200-2000 mm in Hawai’i) and high temperatures (26°C in French Guiana, 22-24°C in Hawaii) (Hijmans, 2015). However, the Australian range is heterogeneous in several environmental factors; on average, sites in QLD are more similar to the native range with respect to aridity, receiving more annual rainfall than sites in the NT and WA (2000-3000 mm in QLD, 400-1000 mm in NT and WA; Fig S1A), and have lower annual mean temperatures (21-24°C in QLD, 24-27°C in NT and WA; Fig S1B) (Bureau of Meteorology, 2018). Heritable phenotypic differences in behavior (J. Gruber, Brown, Whiting, & Shine, 2017), thermal performance (G. K. Kosmala, Brown, Christian, Hudson, & Shine, 2018), morphology (C. M. Hudson, G. P. Brown, & R. Shine, 2016; Hudson, Brown, Stuart, & Shine, 2018), and immune function (Brown, Phillips, Dubey, & Shine, 2015c) have been documented in cane toads at different localities across the Australian range. Nonetheless, surveys have reported low levels of genetic diversity, based on microsatellites (Leblois, Rousset, Tikel, Moritz, & Estoup, 2000) and MHC (Lillie, Shine, & Belov, 2014), and only one mitochondrial haplotype in the *ND3* gene of 31 individuals sequenced from Hawai’i and Australia (Slade & Moritz, 1998).

**Fig 1.**
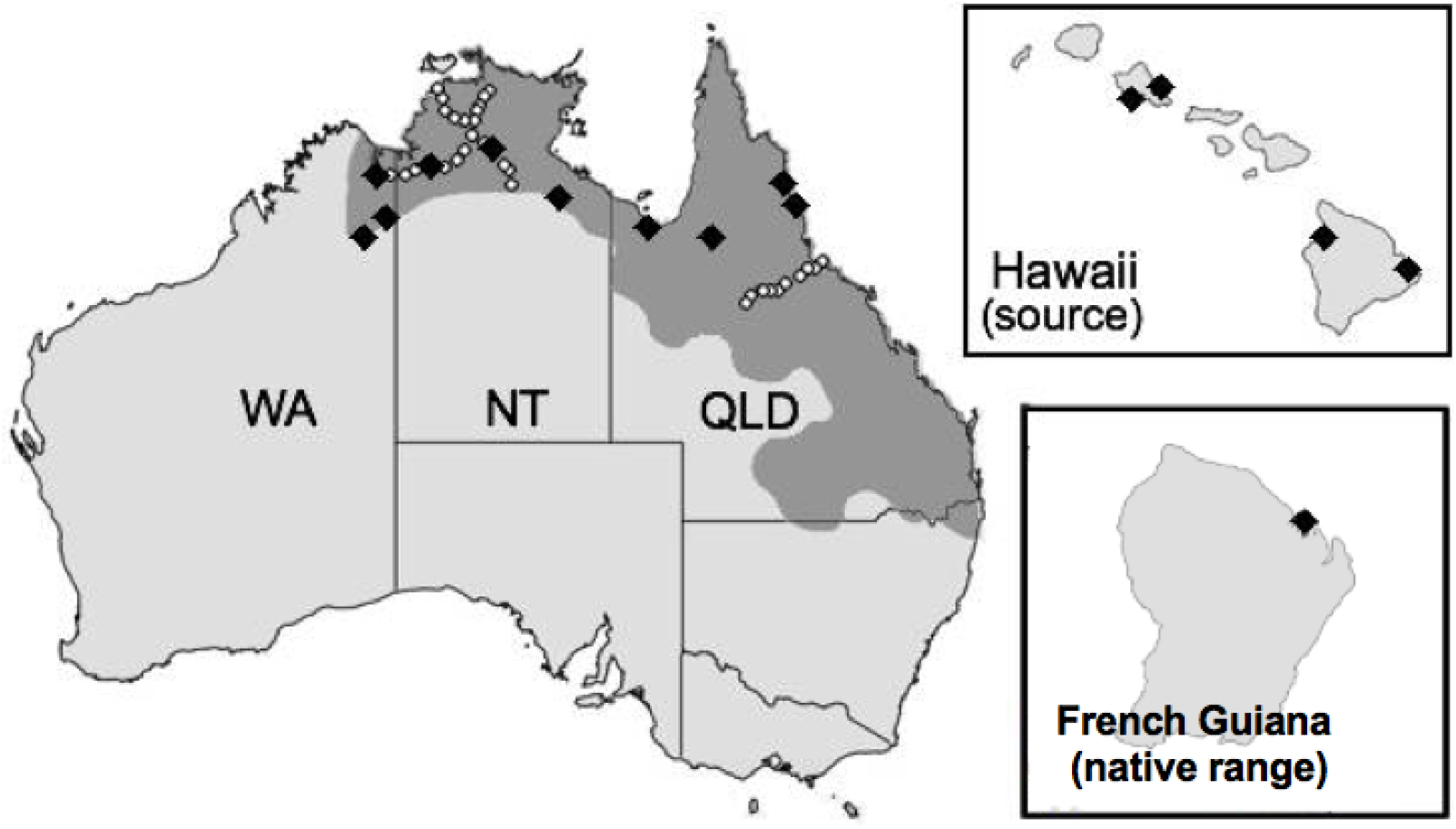
Map of two invaded territories (Australia and Hawai’i) and the native range (French Guiana) of the cane toad (*Rhinella marina*). The shaded region represents the toad’s current Australian range, which is continuing to expand. Black diamonds in French Guiana, Hawai’i, and Australia indicate collection sites for our RNA-Seq experiment. White circles in Australia indicate collection sites for the RADSeq experiment from which we downloaded data.

Here, we assessed genetic diversity and population structure using single nucleotide polymorphisms (SNPs) identified in RNA-Seq data from samples originating from French Guiana (the “native range”), Hawai’i (the “source”), QLD (the “range core”), NT (“intermediate” areas), and WA (the “invasion front”) to test whether the genetic paradox is evident in the cane toad invasion, and to investigate the evolutionary dynamics during introduction. We predicted high divergence between native and invasive populations due to a combination of genetic drift, selection, and spatial sorting, but little genetic differentiation within invasion phases due to a putative lack of standing genetic diversity. Because of the increase in aridity at intermediate areas and the invasion front, we predicted that loci involved in thermal tolerance would be under selection, which may underlie the success that toads have had dispersing through these areas.

RNA-Seq data provide genome-wide information, but are limited to transcribed regions of the genome (Davey & Blaxter, 2011). RNA-Seq data may therefore contain variants that exhibit strong signals for both demography and selection. Conversely, reduced representation sequencing approaches such as restriction site-associated DNA sequencing (RADSeq) provide genome-wide information on a subset of all coding and non-coding sequences (only SNPs near restriction enzyme cleavage sites are detected); because non-coding sequences make up most of the genome, SNPs detected using this technique are more likely to represent neutral variation (Wang, Gerstein, & Snyder, 2009). To focus on how selection may shape genomic diversity in invasive toads, we compared the results from our RNA-Seq data to those from a publicly available RADSeq dataset (Trumbo et al., 2016) that we reanalyzed.

## Materials and Methods

### Sample collection, RNA extraction, and sequencing

We collected samples from French Guiana, the Hawai’ian Islands and across Australia (Fig 1; Table S1). We excised whole spleen tissue (sample sizes in Table S1A) and whole brain tissue (sample sizes in Table S1B) from female toads immediately after euthanasia, and preserved samples in RNAlater (QIAGEN, USA) at −20°C for a maximum of one month, after which they were drained and transferred to a −80°C freezer for long-term storage. We conducted RNA extractions using the RNeasy Lipid Tissue Mini Kit (QIAGEN, USA) following the manufacturer’s instructions, with an additional genomic DNA removal step using on-column RNase-free DNase treatment (QIAGEN, USA). We quantified the total RNA extracted using a Qubit RNA HS assay on a Qubit 3.0 fluorometer (Life Technologies, USA). Extracts were stored at −80°C until sequencing (Macrogen Inc., ROK). mRNA libraries were constructed using the TruSeq mRNA v2 sample kit (Illumina Inc., USA), which included a 300bp selection step. In total, we sequenced 118 individually barcoded samples (46 spleen and 72 brain) across 5 lanes of Illumina HiSeq 2500. Capture of mRNA was performed using the oligo dT method, and size selection parameter choices were made according to the HiSeq2500 manufacturer’s protocol. Overall, this generated 709 million (spleen) and 1.74 billion (brain) paired-end 2 × 125-bp reads. Raw sequence reads are available as FASTQ files in the NCBI short read archive (SRA) under the BioProject Accession PRJNA510261 (spleen data from French Guiana and Hawai’i), PRJNA395127 (spleen data from Australia), and PRJNA479937 (all brain data).

### Data pre-processing and alignment

First, we examined per base raw sequence read quality (Phred scores) and GC content, and checked for the presence of adapter sequences for each sample using FastQC v0.11.5 (S. Andrews, 2010). We then processed raw reads (FASTQ format) from each sample with Trimmomatic v0.35 (Bolger, Lohse, & Usadel, 2014), using the following parameters: ILLUMINACLIP:TruSeq3-PE.fa:2:30:10:4 SLIDINGWINDOW:5:30 AVGQUAL:30 MINLEN:36. This removed any adaptor sequences, trimmed any set of five contiguous bases with an average Phred score below 30, and removed any read with an average Phred score below 30 or sequence length below 36 bp.

As a reference, we used the annotated *R. marina* transcriptome (Richardson et al., 2018), which was constructed from brain, spleen, muscle, liver, ovary, testes, and tadpole tissues. We conducted per sample alignments of reads (FASTQ files) to the reference using STAR v2.5.0a (Dobin et al., 2013) in basic two-pass mode with default parameters, a runRNGseed of 777, and specifying binary alignment map (BAM) alignment outputs. As STAR-generated BAM files lack read groups (identifiers for reads that specify the individual that they come from and the platform that was used to sequence them), we added them to our BAM files using the AddOrReplaceReadGroups tool in Picard Tools (Institute, 2018). To avoid making incorrect variant calls, we removed duplicate reads (reads that map to the same position of the same gene in the same individual) using the MarkDuplicates tool in Picard Tools. To split reads containing an N (region of the reference that is skipped in a read) into individual exon segments, we used the SplitNCigarReads tool in the Genome Analysis Toolkit (GATK) v3.8.0 (McKenna et al., 2010).

### Variant calling and filtering

To call SNPs and insertion-deletions (indels), we used the HaplotypeCaller tool in GATK (McKenna et al., 2010) on our alignment (BAM) files. This tool works by identifying regions of the reference that show evidence of variation (called ‘active regions’). Variants required a minimum phred-scaled confidence of 20 to be called (marked as passing the quality filters) and emitted (reported in the output), using the stand_call_conf and stand_emit_conf options.

To improve accuracy of SNP-calling, we performed this in ‘GVCF’ mode (which generates one intermediate genomic variant call format file per sample). By using this mode, we avoided missing SNPs at loci that match the reference in some but not all individuals. We then used the GenotypeGVCFs tool to merge the GVCF files, re-calculate genotype likelihoods at each SNP locus across all individuals, and re-genotype and re-annotate all SNP loci. The results were written to one merged VCF file. Although we initially genotyped all spleens and brains together, we discovered during downstream analyses that there was an effect of tissue type on population assignment – even from the same individuals, and using only SNPs from transcripts expressed in both tissues, spleen and brain samples were assigned to separate populations). Thus, we genotyped spleens and brains separately, resulting in two merged VCF files, and subsequently kept these separate for all downstream analyses. We retained both datasets because each provides a unique benefit: the spleen dataset includes native range samples, but the brain dataset has more extensive sampling of the invasive populations.

Rather than following a random distribution, some SNPs are clustered. Clustered SNPs may not be independent, and are thus filtered. We used the VariantFiltration tool to identify and filter ‘clusters’ (sets of 3 SNPs that appear within a window of 35 bases) in each of our merged VCF files. We also used this tool to filter variants with QualByDepth (QD; variant confidence divided by the unfiltered depth of non-reference samples) less than 2.0, depth of coverage (DP) less than 20.0, and allele frequency less than 0.05. We then subset our VCF files to include only the variants that passed the filters we set in the VariantFiltration step using the SelectVariants tool. This resulted in 803,489 SNPs from spleen data and 818,536 SNPs from brain data. We used bcftools (Li et al., 2009) to further filter the VCF files, only retaining biallelic SNPs. We examined the results of filtering for minimum minor allele frequency (min MAF) thresholds of 0.01 or 0.05, and several missing data tolerance (MDT) thresholds (the maximum percentage of individuals in the dataset in which a genotype for a locus can be absent without that locus being filtered out). Our population structure results were consistent across min MAF and MDT thresholds, so we ultimately chose to filter our data at min MAF = 0.05 and MDT = 0% (no missing data tolerated) because some downstream analyses cannot handle missing data. These filtering steps reduced the number of SNPs to 65,195 in spleen data and 35,842 in brain data (Table S1A-B).

### Inference of population structure

We used PLINK (Purcell et al., 2007) to convert our VCF files to the Browser Extensible Data (BED) format, which is readable to fastStructure. We then used fastStructure (Raj, Stephens, & Pritchard, 2014) to infer population structure using a variational Bayesian framework for calculating posterior distributions, and to identify the number of genetic clusters in our dataset (K) using heuristic scores (Raj et al., 2014). We ran the structure.py ten times each (K= 1 to 10). We then took the resulting meanQ files from fastStructure and plotted them using the pophelper package (Francis, 2016) in R (Team, 2016).

In addition to using fastStructure for population assignment, we also performed a Redundancy Analysis (RDA) using the vegan package (Oksanen et al., 2018) in R, which fits genetic and environmental data to a multivariate linear regression, and then performs a principal component analysis (PCA) on the fitted values. RDA visualizes both population structure and the effects that environmental variables may have in shaping it. To do this, we downloaded climatic data on French Guiana, Hawai’i, and Australia from the Bioclim database (Hijmans, Cameron, Parra, Jones, & Jarvis, 2005) using the raster package (Hijmans, 2015). Because these areas vary in aridity, we downloaded data on rainfall and temperature; these data are averages of annual statistics over the period of 1970 to 2000. Specifically, we downloaded: annual mean temperature, maximum temperature of the warmest month, minimum temperature of the coldest month, annual precipitation, precipitation of the wettest quarter, and precipitation of the driest quarter. We then used the vcfR package (Knaus & Grünwald, 2017) to convert our VCF files to the GENLIGHT format, which is readable to the vegan package.

### Identification of candidate loci under selection

Loci that are under natural selection may have abnormally high F_ST_ values, causing them to be outliers among all other loci in the transcriptome. Bayescan v2.1 (Foll & Gaggiotti, 2008) detects loci with outlier F_ST_s using the multinomial-Dirichlet model. We used PGDSpider (Lischer & Excoffier, 2012) to convert our VCF files to the BAYESCAN format, and then ran Bayescan to detect loci with outlier pairwise F_ST_ values between our three genetic clusters in spleen data (native range toads versus source and core toads versus intermediate and frontal toads) and our two genetic clusters in brain data (source and core toads versus intermediate and frontal toads).

Loci under natural selection also may have allele frequencies that are associated with environmental variables. We used the climatic data that we downloaded from the Bioclim database (Hijmans et al., 2005) using the raster package (Hijmans, 2015) on rainfall during the driest quarter and maximum temperature in the warmest month. We then used the lfmm v2.0 package (Frichot & Francois, 2015) in R to perform a latent factor mixed model (LFMM) to test the association between the allele frequencies of every locus and these two environmental variables. We applied a Benjamini-Hochberg correction to all p-values from the LFMM.

Because these scans reveal thousands of loci putatively under selection, and because outlier tests have high false positive rates (Whitlock & Lotterhos, 2015), we took a more conservative approach by cross-matching our list of F_ST_ outliers with each of our lists of environmentally-associated loci. For individual investigation, we only looked at loci with both an outlier FST value and an association with a putatively significant environmental variable.

### Evaluation of genetic differentiation and diversity

To quantify levels of genetic differentiation and diversity, we computed basic statistics in the hierfstat (Goudet, 2005) and diveRsity (Keenan, McGinnity, Cross, Crozier, & Prodohl, 2013) packages in R (Team, 2016). We first used PGDSpider v2.1.1.3 (Lischer & Excoffier, 2012) to convert our VCF files to the FSTAT and GENEPOP formats, which are readable to hierfstat and diveRsity, respectively. We then used these packages to calculate measures of genetic differentiation, including global FST and pairwise FST (by genetic cluster, invasion phase, and collection site), and genetic diversity, such as expected heterozygosity (He) and rarefied allelic richness (AR). We also computed Shannon’s Information Index (SI) with the dartR package (B. Gruber, Unmack, Berry, & Georges, 2018). After calculating these measures of diversity across all loci (N=65,195 from spleen data, N=35,842 from brain data), we calculated the same measures in loci putatively under selection: those with outlier F_ST_ values (N=648 from spleen data, N=203 from brain data) and those associated with environmental variables (N=4179 from spleen data, N=530 from brain data). In spleen data, this allowed us to investigate the hypothesis that genetic diversity is maintained at ecologically relevant traits even if genome-wide diversity is lost. In brain data, this allowed us to examine the effect of natural selection on genetic diversity within Hawai’i and Australia. We used Kruskal-Wallis tests to assess the significance of the differences in genetic diversity.

### Annotation of SNPs

To visualize the types of genomic regions in which our SNPs lie, we used SnpEff (Cingolani et al., 2012). SnpEff analyzes the information in a VCF file, annotates the SNPs, and estimates their effects. We calculated the relative proportions of each type of SNP for both tissue types using the full dataset of SNPs, and our two selection candidate datasets (containing F_ST_ outlier loci associated with an environmental variable). We then used the stats v3.5.0 R package to perform z-tests on the differences in proportions of SNP types between the full and selection candidate datasets. We expected differences in the proportions of certain types of SNPs, such as a lower proportion of synonymous variants in the selection candidates because synonymous mutations do not result in a codon change (and thus are unlikely to change protein function, i.e. phenotype).

### Isolation by distance

To examine the effects of geographic distance on genetic distance across the Australian range, we performed a Mantel test using ade4 v1.7-5 (Thioulouse & Dray, 2007). Because we were focused on isolation by distance through range expansion after introduction (to Australia), samples from Hawai’i and French Guiana were excluded; and because we sampled brain tissue in a greater number of sites within Australia (as opposed to spleen tissue; Table S1A-B), we only performed the Mantel test on brain data. For the geographic distance matrix, we used the dist function in R (Team, 2016) to calculate the Euclidean distances in geographic space between collection sites based on their coordinates. For the genetic distance matrix, we used the collection site-based pairwise F_ST_ values generated from hierfstat. We performed four Mantel tests, using: (1) all loci, (2) loci with outlier F_ST_ values, (3) loci with outlier F_ST_ values and with an environmental association (temperature or rainfall), (4) loci with outlier F_ST_ values but without an environmental association. A significant result across all loci may suggest that the prominent driver of genetic structure is genetic drift. However, clinal variation in allele frequencies may also result from spatial sorting (particularly in loci underlying dispersal ability), or from selection associated with environmental clines; distinguishing these evolutionary processes is difficult. Thus, we performed the additional tests to attempt to separate the effects of genetic drift and spatial sorting. Loci with outlier FST values are candidates for natural selection and spatial sorting; thus, a significant result from only these loci would less likely be caused by genetic drift. Outlier loci can be further separated into those with an environmental association, which are likely candidates for natural selection driven by environmental variables, and those without an environmental association, which are possible candidates for spatial sorting. If spatial sorting is indeed influencing genetic differentiation, then we may expect to see different patterns between these two groups of loci (e.g. linearity only in the outlier loci without an environmental association).

### Acquisition, processing, and analysis of RADSeq dataset

In addition to our own RNA-Seq dataset, we downloaded a publicly available RADSeq dataset (Trumbo et al., 2016) to evaluate population structure in Australian cane toads. Unlike RNA-Seq data, which consists of coding sequences only (Wang et al., 2009), RADSeq data include SNPs from across the entire genome, but only around sites that are cleaved by a selected set of restriction enzymes (Davey & Blaxter, 2011). This method allows us to contrast the results of assessing loci potentially under ‘tighter’ selection (e.g., codon bias) with those primarily constituting neutral variation (RADSeq). Individuals in the RADSeq dataset were collected from QLD (range core towards intermediate areas, N=179) and the border of NT/WA (intermediate areas towards the invasion front, N=441; Fig 1). However, samples were lacking from a large part of the range between those regions; thus, identifying geographic points of differentiation between genetic clusters may be unlikely using this dataset. Furthermore, although there were many collection sites within these regions, metadata with coordinates (or even location names) were unavailable. We downloaded raw 100-bp reads from NCBI short read archive (SRA) under the BioProject Accession PRJNA328156. We then converted SRA files to the FASTQ format using the fastq-dump tool in the SRA toolkit.

We used Stacks v2.0 (Catchen, Hohenlohe, Bassham, Amores, & Cresko, 2013) for all RADSeq data processing. First, we used the process_radtags program to remove low quality reads from the FASTQ files. Next, we used the denovo_map pipeline to perform a *de novo* assembly for each sample, align matching DNA regions across samples (called ‘stacks’), and call SNPs using a maximum likelihood framework. We allowed a maximum of five base mismatches between stacks within an individual and three base mismatches between stacks between individuals. This resulted in 499,623 SNPs. We then filtered the results using the populations program, using one random SNP per locus and four combinations of parameter choices (min MAF=0.05, MDT = 0.5; min MAF = 0.05, MDT = 0.2; min MAF = 0.01, MDT = 0.5; min MAF = 0.01, MDT = 0.2). Structure results were consistent across min MAF values, but not MDT values: two distinct populations were only detected using the more stringent MDT choice. Because population structure in this dataset is relatively low, and because it is appropriate to use more stringent filtering choices when analyzing datasets with low population structure (Selechnik et al., unpublished; Chapter 2), we ultimately chose to filter this dataset with min MAF = 0.05 and MDT = 0.2. Furthermore, using these filtering choices produced results that were most consistent with those reported by Trumbo et al. (2016) and with those of our RNA-Seq dataset. Filtering reduced the number of SNPs to 8,296. The results were written to a VCF file and a STRUCTURE file (which could be read into fastStructure).

We ran fastStructure using the same methodology as we did for our RNA-Seq dataset. In addition to the model-based fastStructure, we also performed a discriminant analysis of principal components (DAPC) on this dataset using adegenet (Jombart & Ahmed, 2011). DAPC is a multivariate approach that identifies the number of genetic clusters using K-means of principal components and a Bayesian framework. We also converted the VCF to the FSTAT and GENEPOP formats and computed the same basic statistics for the RADSeq dataset as we did for our RNA-Seq datasets using hierfstat and diveRsity. Because collection site coordinate metadata were unavailable for the RADSeq dataset, we did not perform a Mantel test using these data.

## Results

### Population structure across the Hawai’ian-Australian invasion

We found three genetic clusters using spleen data (Fig 2A): French Guiana (native range) formed its own genetic cluster, Hawai’i (source) clustered with QLD (core), and NT (intermediate) clustered with WA (invasion front). In our brain dataset, population differentiation seemed to align with environmental barriers (Fig 2B-C): Hawai’i (source) clustered with coastal QLD (core), whereas inland QLD/NT (intermediate) clustered with WA (invasion front). Inference of substructure within these two genetic clusters revealed that source and core toads may further differentiate into two separate groups (Fig 2D); however, intermediate and frontal toads remain one genetic cluster (Fig 2E).

**Fig 2.**
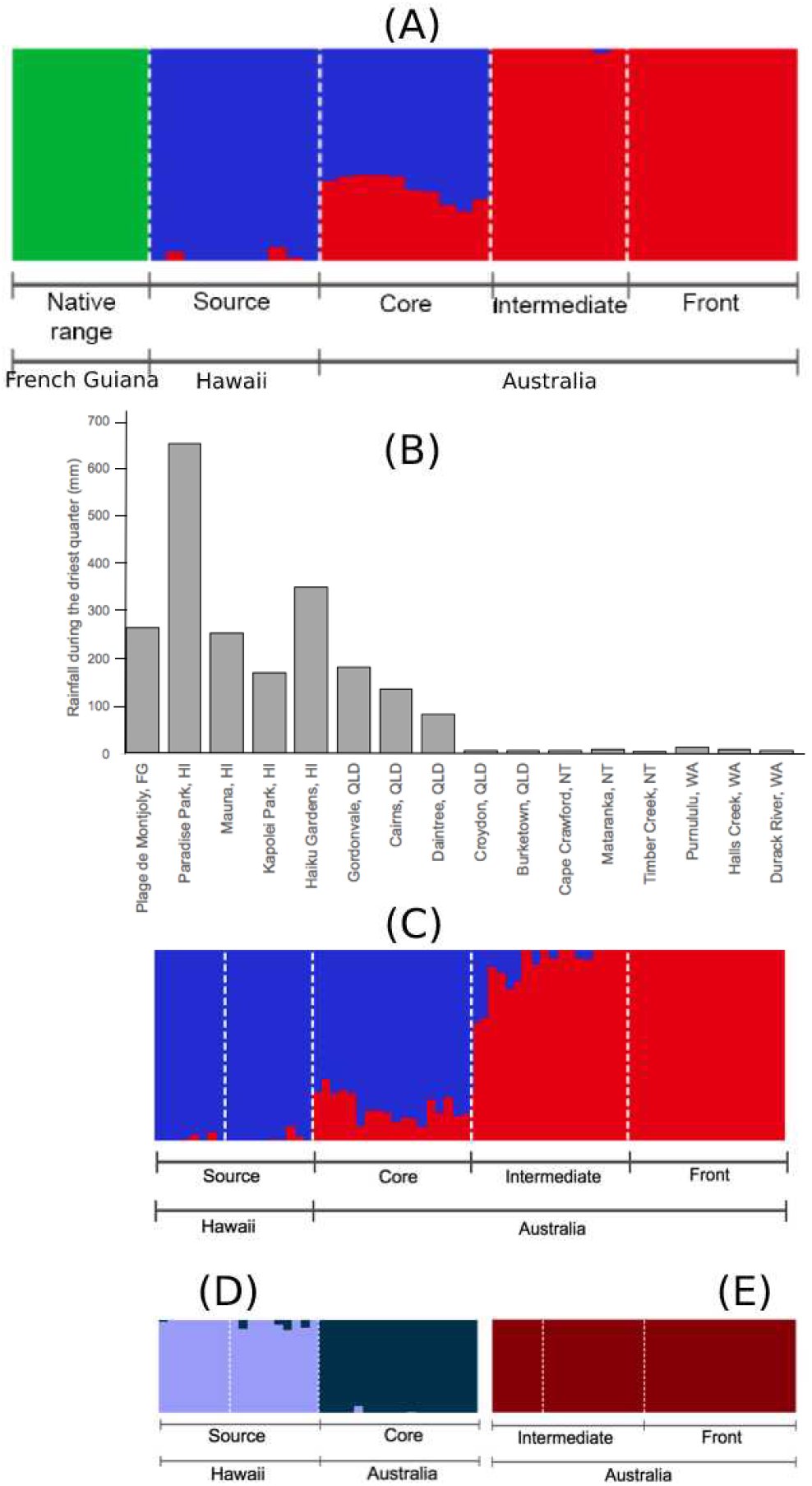
(A) Population structure of cane toads from French Guiana, Hawai’i, and Australia, as inferred by SNPs from spleen RNA-Seq data. (B) Rainfall during the driest quarter in each location across collection sites for invasive cane toads in French Guiana, Hawaii, and Australia. A similar environmental barrier occurs in temperature (Fig S1). (C) Population structure of invasive cane toads from Hawai’i and Australia, as inferred by SNPs from brain RNA-Seq data. (D) Substructure within toads from Hawai’i (HI) and Queensland (QLD) as inferred by SNPs from brain RNA-Seq data. (E) Substructure within toads from Northern Territory (NT) and Western Australia (WA) as inferred by SNPs from brain RNA-Seq data.

RDA was mostly consistent with fastStructure for both spleen (Fig 3A) and brain (Fig 3B) data, except for the differentiation of source toads and core toads as two separate genetic clusters. It also suggested that: (1) Source toads may have diverged from toads of other clusters due to selection from rainfall during the driest quarter. (2) Core toads may have diverged from toads of other clusters due to selection from rainfall during the wettest quarter, mean annual rainfall, and minimum temperature during the coolest month. (3) Intermediate and frontal toads may have diverged from toads of other clusters due to selection from maximum temperature during the hottest month and mean annual temperature.

**Fig 3.**
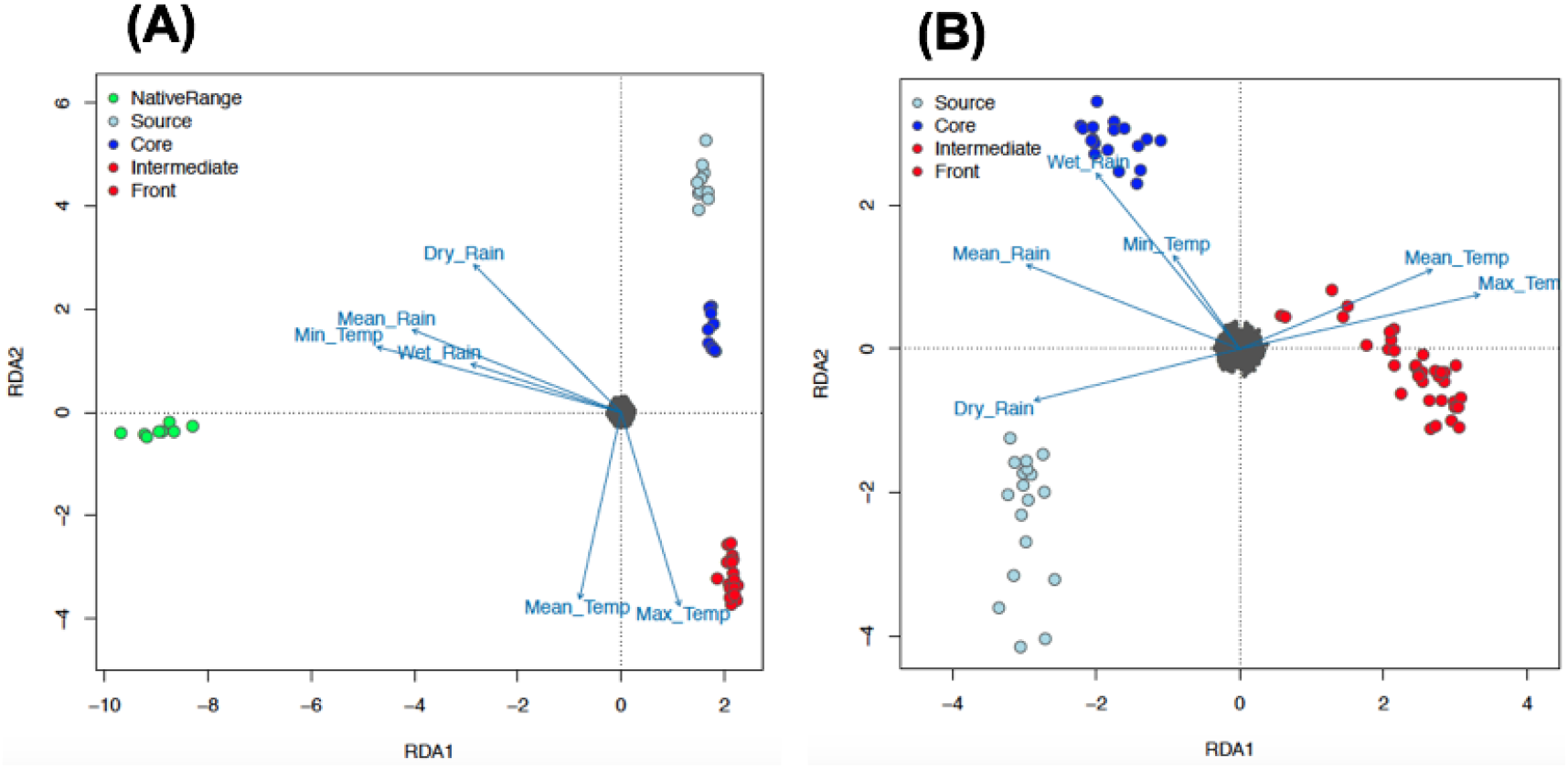
(A) Relationship between environmental variables and population structure of cane toads from French Guiana, Hawai’i, and Australia, as inferred by Redundancy Analysis (RDA) on SNPs from spleen RNA-Seq data. Environmental data include: annual mean temperature (Mean Temp), maximum temperature of the warmest month (Max Temp), minimum temperature of the coldest month (Min Temp), annual precipitation (Mean Rain), precipitation of the wettest quarter (Wet Rain), and precipitation of the driest quarter (Dry Rain). (B) Relationship between environmental variables and population structure of invasive cane toads from Hawai’i and Australia, as inferred by RDA on SNPs from brain RNA-Seq data.

### Tests for selection

Out of 65,195 SNPs from spleen data, 648 were identified as F_ST_ outliers, with a mean global F_ST_ of 0.89. Mean pairwise F_ST_ values across outlier loci were 0.93 between the native and source/core populations, 0.97 between the native and intermediate/front populations, and 0.34 between the source/core and intermediate/front populations. Also from the spleen dataset, 3088 SNPs were associated with maximum temperature during the warmest month and 1813 were associated with rainfall during the driest quarter. Of these loci, 772 were associated with both environmental variables. We focused on SNPs that were detected both as F_ST_ outliers and environmental correlates: there were 64 outlier SNPs associated with maximum temperature and 47 outlier SNPs associated with rainfall in the driest quarter (including 18 outliers associated with both environmental variables).

Most of the 64 outlier SNPs associated with maximum temperature during the hottest month (Table S2A) were in transcripts with functions such as metabolism, transcription regulation, and immune function. There were also two SNPs in *HSP4*, a gene encoding heat shock protein (HSP) 4 (70 kDa), which is involved in thermal tolerance (on exposure to heat stress) and protein folding and unfolding (Consortium, 2017). Most of the 47 outlier SNPs associated with rainfall during the driest quarter (Table S3A) were in transcripts with functions such as cell signaling and immune function.

Out of 35,842 SNPs from brain data, 203 were identified as F_ST_ outliers, with a mean global F_ST_ of 0.34. Mean pairwise F_ST_ was 0.50 between the source/core and intermediate/front populations. Also from the brain dataset, 345 SNPs were associated with maximum temperature during the warmest month and 194 were associated with rainfall during the driest quarter. Nine of these loci were associated with both environmental variables. Of SNPs that were detected both as F_ST_ outliers and environmental correlates, there were 38 outlier SNPs associated with maximum temperature and 33 outlier SNPs associated with rainfall in the driest quarter; no outliers were associated with both environmental variables.

Most of the 38 outlier SNPs associated with maximum temperature during the hottest month (Table S2B) were in genes involved in cell signaling, mitochondrial processes (metabolism), or gene expression (Consortium, 2017). Similarly to the spleen data, there was one SNP from brain data in *HSP4*. Another SNP was in *ARNT2*, a gene encoding aryl hydrocarbon receptor nuclear translocator 2, a transcription factor involved in the hypothalamic pituitary adrenal (HPA) axis and visual and renal function (Consortium, 2017). Of the 33 outlier SNPs from brain data correlated with rainfall during the driest quarter (Table S3B), 19 were in *MAGED2*, a gene encoding melanoma-associated antigen D2, which is involved in renal sodium ion absorption (Consortium, 2017). Four were in *STXBP1*, a gene encoding syntaxin-binding protein 1, which is involved in platelet aggregation and vesicle fusion (Consortium, 2017). The rest were in genes generally involved in gene expression and cell signaling (Consortium, 2017).

### Evaluation of genetic differentiation, diversity, and isolation by distance

Our spleen data show strong divergence between the native and source/core populations (pairwise F_ST_ = 0.29), and even stronger divergences between native and intermediate/frontal populations (pairwise F_ST_ = 0.32). Differentiation between the source/core and intermediate/frontal populations was much lower (pairwise F_ST_ = 0.08). Overall divergence among natives and invaders was moderately high (global F_ST_ = 0.18)

In brain data, genetic differentiation between the source/core and intermediate/frontal populations also was low (pairwise F_ST_ = 0.07). Differentiation between source and core toads was lower when calculated with brain data as compared to spleen (pairwise FST = 0.04). Generally, pairwise F_ST_ by invasion phase (Table S4A) and collection site (Table S4B) calculated from brain data revealed that toads from areas that are more closely linked by invasion history are less strongly differentiated. Because the brain data did not include native range samples, overall divergence was estimated to be much lower than it was in the spleen data (global F_ST_ = 0.04).

Diversity statistics from spleen data suggest that the native range population is only slightly more genetically diverse than the source/core population (native: He = 0.27, AR = 1.74, SI = 0.40; source/core: He = 0.26, AR = 1.74, SI = 0.40; *p* = 5E-4 for He) but both are more diverse than the intermediate/frontal population (He = 0.23, AR = 1.68, SI = 0.36; *p* < 2E-16 for all measures). However, we also estimated these measures by region (Table 1) rather than by genetic cluster to detect changes in genetic diversity at a finer scale; these calculations reveal a more obvious decline in genetic diversity from the native population to all invasive collection sites.

**Table 1.**
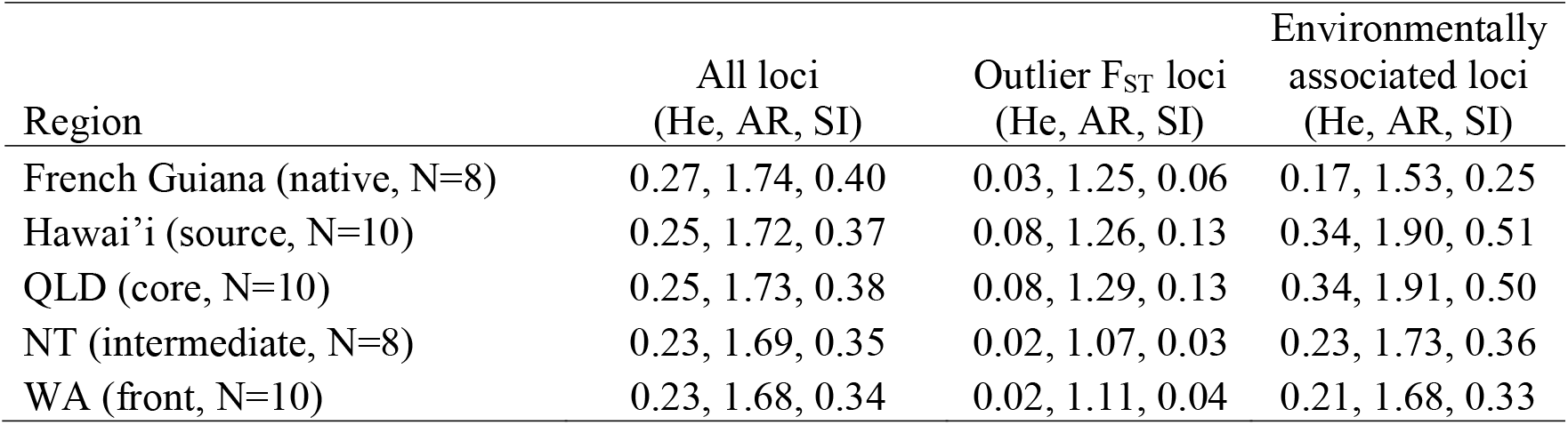
Expected heterozygosity (He), allelic richness (AR), and Shannon’s Information Index (SI) estimated using SNPs from spleen RNA-Seq data on native and invasive populations of cane toads *(Rhinella marina)* at regions across French Guiana, Hawai’i, and Australia. QLD = Queensland, NT = Northern Territory, WA = Western Australia.

Despite this reduction in genome-wide genetic diversity, subsets of loci putatively under selection in the spleen dataset showed no loss of diversity during introduction (Table 1). Here there was an increase in diversity in the source/core population as compared to native samples (native: He = 0.03, AR = 1.25, SI = 0.06; source/core: He = 0.09, AR = 1.37, SI = 0.14; *p* < 2E-16 for all measures). At these same loci, diversity was lowest in the intermediate/front population (He = 0.02, AR = 1.14, SI = 0.04). These trends are also shown in environmentally-associated loci (native: He = 0.17, AR = 1.53, SI = 0.25; source/core: He = 0.37, AR = 1.99, SI = 0.55; intermediate/front: He = 0.23, AR = 1.81, SI = 0.35).

Genome-wide diversity estimates using brain data show similar patterns to spleen data (source/core: He = 0.34, AR = 2.00, SI = 0.51; intermediate/front: He = 0.31, AR = 1.97, SI = 0.47; *p* < 2E-16 for comparisons of all measures; Table 2). This reduction is greater in loci with outlier F_ST_ values (source/core: He = 0.38, AR = 2.00, SI = 0.55; intermediate/front: He = 0.15, AR = 1.70, SI = 0.24; *p* < 2E-16 for comparisons of all measures) and in environmentally-associated loci (source/core: He = 0.35, AR = 2.00, SI = 0.53; intermediate/front: He = 0.08, AR = 1.57, SI = 0.15; *p* < 2E-16 for comparisons of all measures).

**Table 2.** Expected heterozygosity (He), allelic richness (AR), and Shannon’s Information Index (SI) estimated using SNPs from brain RNA-Seq data on native and invasive populations of cane toads (*Rhinella marina*) at regions across French Guiana, Hawai’i, and Australia. QLD = Queensland, NT = Northern Territory, WA = Western Australia.

| Region | All loci<br>(He, AR, SI) | Outlier $F_{ST}$ loci<br>(He, AR, SI) | Environmentally<br>associated loci<br>(He, AR, SI) |
| --- | --- | --- | --- |
| Hawai'i (source, N=18) | 0.33, 1.97, 0.49 | 0.35, 1.87, 0.51 | 0.41, 2.00, 0.60 |
| QLD (core, N=18) | 0.33, 1.98, 0.50 | 0.39, 2.00, 0.56 | 0.24, 1.92, 0.38 |
| NT (intermediate, N=18) | 0.31, 1.95, 0.47 | 0.17, 1.66, 0.28 | 0.09, 1.52, 0.16 |
| WA (front, N=18) | 0.29, 1.93, 0.45 | 0.12, 1.49, 0.19 | 0.07, 1.46, 0.12 |

Geographic distance in collection sites and pairwise F_ST_ values by collection site (as estimated with brain data) were significantly associated across all four of our Mantel tests using: (1) all loci (*p* = 1E-3, *r* = 0.76; Fig 4A); (2) all loci with outlier F_ST_ values (*p* = 1E-3, *r* = 0.75; Fig 4B); (3) only loci with outlier F_ST_ values and an environmental association (*p* = 1E-3, *r* = 0.62; Fig 4C); (4) only loci with outlier F_ST_ values but without an environmental association (*p* = 1E-3, *r* = 0.77; Fig 4D).

**Fig 4.**
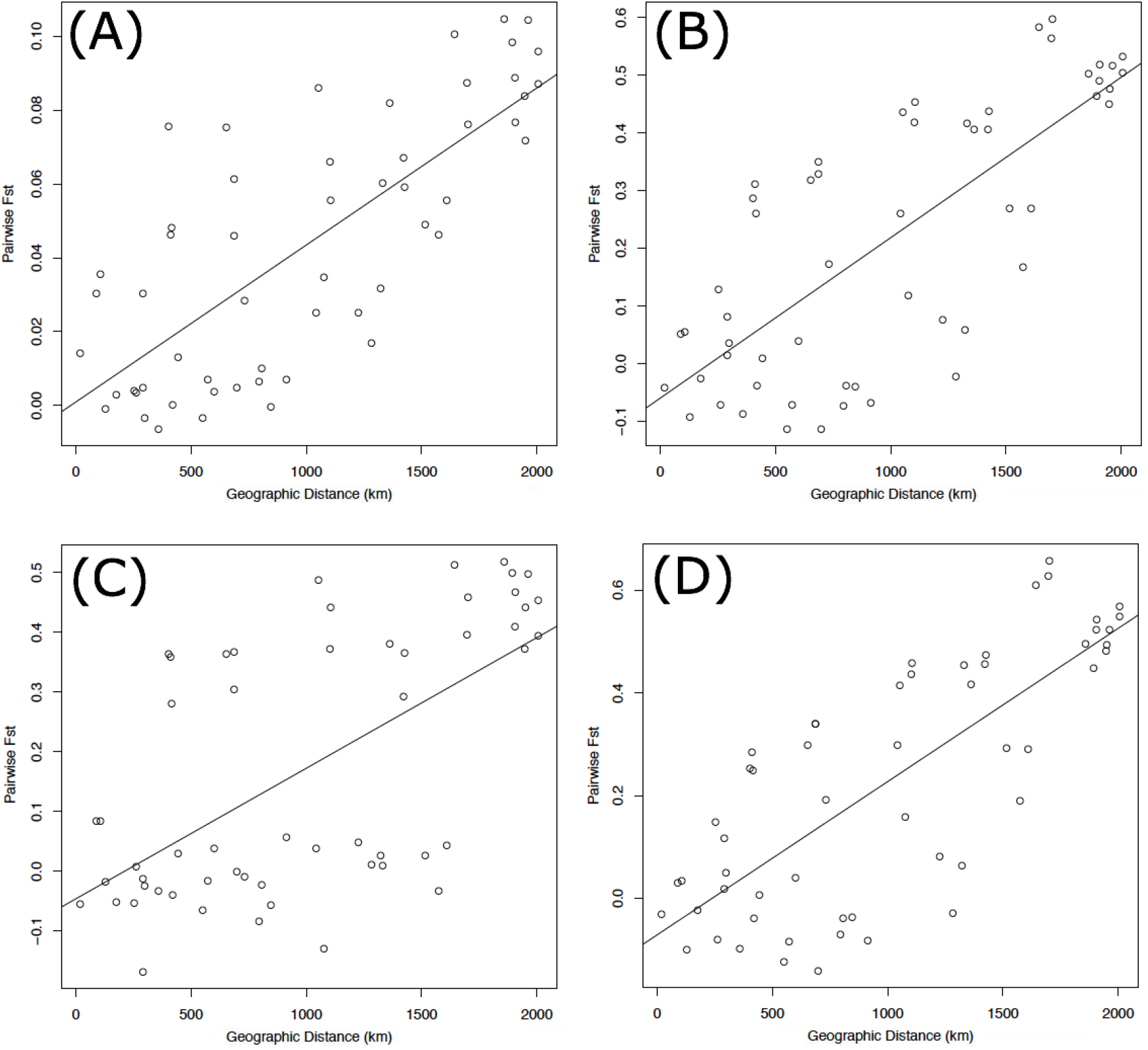
Relationship between geographic distance (km) and genetic distance (pairwise F_ST_) between cane toads collected across their invasive Australian range (Fig 1) as inferred by brain RNA-Seq data. (A) represents the full dataset of loci (*p* = 1E-3, *r* = 0.76). (B) represents all F_ST_ outliers (*p* = 1E-3, *r* = 0.75). (C) represents only F_ST_ outliers with an association with either rainfall or temperature (*p* = 1E-3, *r* = 0.62). (D) represents only F_ST_ outliers without an association with either rainfall or temperature (*p* = 1E-3, *r* = 0.77).

### SNP Annotations

Annotation of SNPs revealed that most of our loci were either missense, synonymous, or found in the 3’ untranslated region (UTR). A small percentage were found in the 5’ UTR, and the remaining loci involved the loss or gain of stop or start codons (Figs S2-S3). In the spleen data, the temperature-associated outlier FST subset had a significantly higher proportion of synonymous variants (*p* = 8E-3) than the full set of SNPs. The outlier loci within *HSP4* were both synonymous variants. However, there were no significant differences in proportions between the rainfall-associated outlier F_ST_ subset and the full set of SNPs.

In the brain data, there were no significant differences in proportions of SNP variant types between the temperature-associated outlier FST subset and the full set. The locus within *HSP4* was a 3’ UTR variant, and the locus within *ARNT2* was synonymous. The rainfall-associated outlier F_ST_ subset had a significantly higher proportion of 3’ UTR variants (*p* = 5E-3) and lower proportion of missense variants (*p* = 0.04) than did the full set. Eighteen of the nineteen loci within *MAGED2* were 3’ UTR variants (the last was missense), and two of the four loci within *STXBP1* were missense (the other two were synonymous).

### Population structure in Australian toads from the RADSeq experiment

Analysis of the RADSeq dataset through fastStructure and DAPC also showed two genetic clusters within Australia (Fig 5A-B): QLD (core towards intermediate areas) in the first group, and the border of NT/WA (intermediate areas towards the invasion front) in the second group. There was little differentiation between these two groups (pairwise F_ST_ = 0.09) or overall (global F_ST_ = 0.04). We found no evidence of significant substructure within either of the two genetic clusters. The two groups were equal in allelic richness (AR = 2.00 for both groups), but expected heterozygosity was higher in core toads than in intermediate/frontal toads (core: He = 0.50; intermediate/front: He = 0.33).

**Fig 5.**
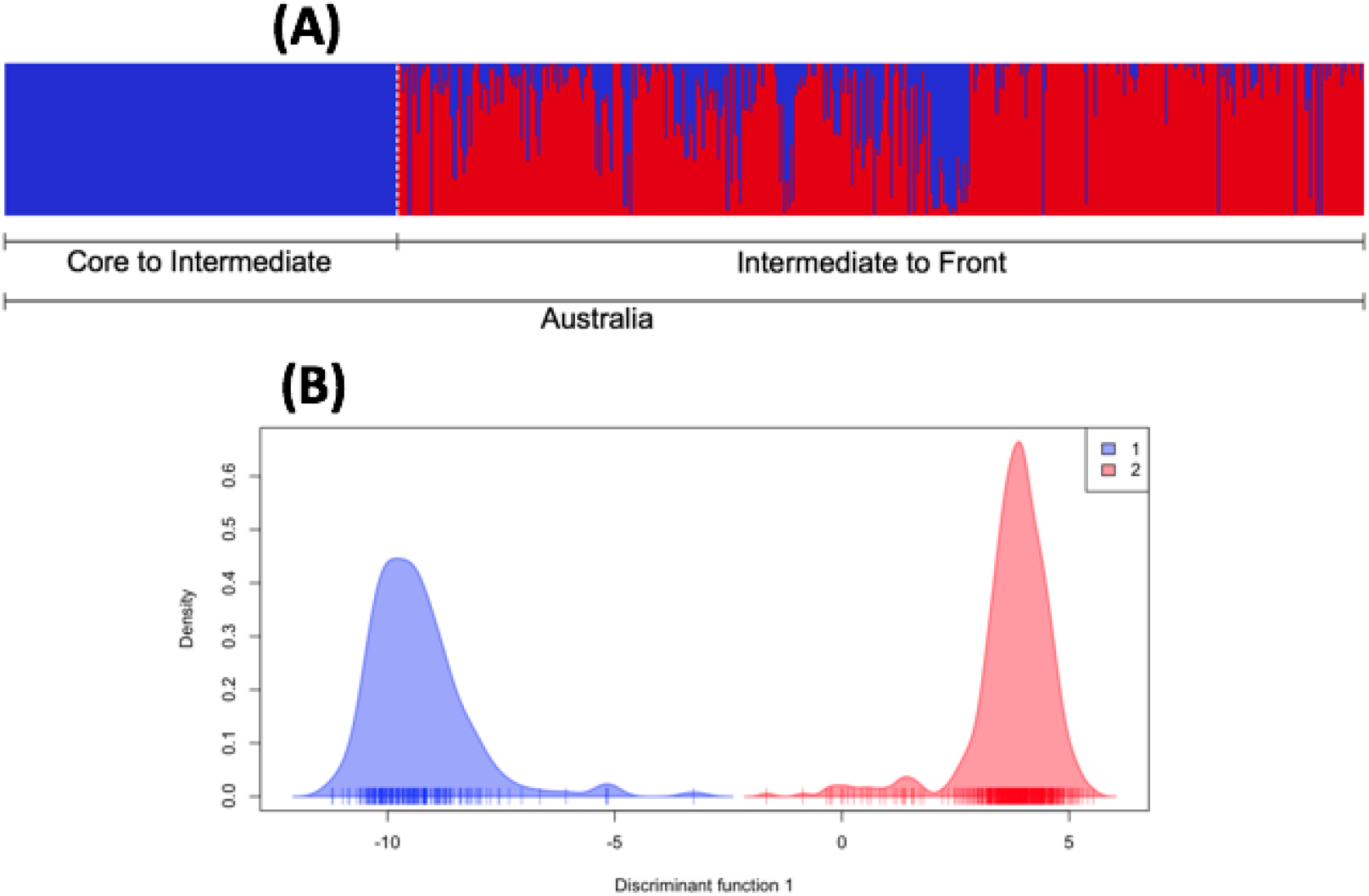
(A) Population structure of cane toads across their invasive range in Australia, as inferred from running fastStructure on SNPs from RADSeq data. Because coordinate metadata for collection sites were unavailable, the samples could not be arranged from east to west along the Australian range. (B) Population structure of cane toads across their invasive range in Australia, as inferred from DAPC on SNPs from RADSeq data.

## Discussion

We predicted that genetic diversity would be significantly lower in invasive populations than in native populations due to genetic bottlenecks occurring during serial introductions of cane toads to Hawai’i and Australia. We also predicted strong divergence between native range and invasive populations due to a combination of genetic drift, selection and spatial sorting. In terms of population structure, we predicted low genetic structure within invasive populations in Hawai’i and Australia resulting from putatively low levels of standing genetic variation. Our analyses show evidence of a reduction in genetic diversity from native to invasive populations across all loci. We found three genetic clusters: (1) native range toads, (2) Hawai’ian source toads and eastern Australian range core toads, and (3) all toads from further west in more recently colonized areas (intermediate and invasion front). Differentiation is much higher between the native population and the invasive populations than between the invasive populations. Loci identified as FST outliers had values greater than 0.9 between native and invasive populations, indicating that many of these loci are likely to be fixed for different alleles in these populations. Genetic structure around the aridity barrier suggest that natural selection is occurring in the western part of the Australian range. However, the presence of isolation by distance across all loci also supports gradual genetic drift across invasion phases; that this relationship is strengthened in the subset of F_ST_ outliers without environmental associations may indicate the presence of spatial sorting. It seems likely that all these evolutionary forces have shaped Australian invasive populations.

Evidence that invasive cane toads fit the genetic paradox paradigm is equivocal. We did identify a reduction in overall genetic diversity, both in allelic richness and evenness (He and SI). Previous studies on mtDNA (Slade & Moritz, 1998) and microsatellite (A. Estoup, Wilson, Sullivan, Cornuet, & Moritz, 2001) data have also suggested losses in genetic diversity from the native range to Hawai’i and Australia. Despite these findings, we did not find this reduction in the subset of loci putatively under selection (loci with outlier F_ST_ values or associations with environmental variables). This supports the prediction that some invasions do not represent a genetic paradox, despite overall loss of genetic diversity and the presence of novel adaptive challenges, because genetic diversity at ecologically relevant traits is maintained by balancing selection (A. Estoup et al., 2016). However, diversity of this subset of loci is not only maintained in the source/core population, it is higher than in the native population. This may reflect admixture from multiple genetically-distinct introductions; the Hawai’ian introduction was sourced from the Puerto Rican population, which included introductions from French Guiana and Guyana (Turvey, 2009). Alternatively, this may represent a genuine paradox; *de novo* mutation rates may have been high enough to restore or even enhance adaptive potential in the source/core population (A. Estoup et al., 2016). Although these toads have been in Hawai’i and Australia for less than 100 years, they were separated from native toads in the early 1800s when toads were first introduced to Caribbean islands. From the source/core population to the intermediate frontal population, however, diversity of loci putatively under selection is highly reduced; we believe this is because of directional selection, as outlined below.

Our two invasive genetic clusters (source/core and intermediate/front) diverge at the transition from coastal to inland areas within Australia, in an area where temperatures become higher and rainfall becomes scarcer, particularly during the dry season (Bureau of Meteorology, 2018). This correlation suggests that these environmental variables may drive population structure and that toads may be adapting to local conditions. Furthermore, RDA indicates selection from temperature and rainfall may drive divergence between invasive populations. Heterogeneous climates across introduced or expanding ranges have previously been linked to genetic differentiation, possibly due to climatically-imposed selection (Leydet et al., 2018; Weber & Schmid, 1998).

Diversity estimates from our brain data suggest a reduction in diversity from the source/core population to the intermediate/frontal population in the full set of loci, but a much greater reduction in diversity in the subset of loci putatively under selection. The source (Hawai’i) and core (QLD) areas represent earlier phases of the invasion and experience similar environmental conditions to those of the native range, whereas intermediate (NT) and invasion front (WA) areas are more arid. Thus, the greater depletion in diversity at loci putatively under selection across the Australian invasion may be due to the effects of directional selection from the harsher climate at more recently colonized areas of the Australian range (C. A. Andrews, 2010; A. Estoup et al., 2016).

Among our loci associated with temperature are several within a gene encoding protein HSP 4 (70 kDa). HSPs protect protein folding during increased temperatures and provide cells with enhanced thermal tolerance (Kiang & Tsokos, 1998), and expression of HSP genes has been shown to underlie adaptive responses to environmental stress (Chen, Feder, & Kang, 2018). Changes in HSP levels in response to thermal environment manipulation vary between native and invasive cane toads, as well as between populations of toads within Australia (G. K. Kosmala, Brown, & Shine, 2018). In our study, the SNPs within *HSP4* included two synonymous variants (spleen data) and a 3’ UTR variant (brain data); 3’ UTR variants are known to alter levels of protein functioning through differential expression of genes (Skeeles, Fleming, Mahler, & Toland, 2013). In the case of HSPs, adaptation may not necessarily mean a change in function, but rather a change in the expression levels of proteins involved in the function, which could be caused by a 3’ UTR variant.

Aryl hydrocarbon receptor nuclear translocator 2 (from the *ARNT2* gene, also associated with temperature), is a transcription factor that specifically acts on genes involved in the HPA axis and visual and renal function. The HPA axis is stimulated by high temperatures (Malmkvist et al., 2009), and the secretion of corticosterone can be modulated adaptively in response to thermal change (Telemeco & Addis, 2014). Elevation of corticosterone secretion in toads increases the rate of desiccation, thus making it maladaptive in arid environments (Jessop, Letnic, Webb, & Dempster, 2013). Corticosterone levels in Brazilian and Australian toads in response to manipulation of thermal environment display similar patterns to those of HSPs (G. K. Kosmala, Brown, & Shine, 2018). Selection on loci potentially affecting corticosterone secretion by affecting the HPA is consistent with these results, although the SNP we investigated in *ARNT2* is synonymous.

More than half (19 of 33) of the loci under selection associated with rainfall (in brain data) were found in *MAGED2* (encodes melanoma-associated antigen D2). This protein regulates the expression and localization of two co-transporters that facilitate renal sodium ion absorption, preventing excess loss of water and solutes via reabsorption through the kidneys (Greger, 2000). Similarly to the SNP within *HSP4*, 18 of 19 SNPs within *MAGED2* were 3’ UTR variants, which can affect gene expression (Skeeles et al., 2013). The high number of SNPs found in the 3’ UTR of *MAGED2* is likely responsible for the difference in proportions of variant types between the full dataset and the rainfall-associated outlier dataset. Four SNPs (including missence variants) were found in *STXBP1* (encodes syntaxin-binding protein 1, which is involved in vesicle fusion and blood clotting). Dehydration is known to increase blood clotting rate (El-Sabban, Fahim, Al Homsi, & Singh, 1996), and excessive blood clotting can be harmful to the heart, brain, and limbs (PubmedHealth, 2014). Evolved changes in the rates of renal sodium absorption and blood clotting may allow toads from intermediate and invasion front areas to survive in drier environments. The remaining SNPs associated with rainfall or temperature were in genes generally involved in gene expression or signal transduction; these genes may facilitate expression of (or signaling to) the genes we have discussed, or may have independent functions in cellular maintenance.

Invasive species have been shown to rapidly adapt phenotypically to heterogeneous climatic conditions, enabling range expansion (Colautti & Barrett, 2013). Some invaders tolerate higher temperatures (Braby & Somero, 2006) and water loss (Godoy, Lemos-Filho, & Valladares, 2011) better than do related indigenous taxa (Zerebecki & Sorte, 2011). There is evidence of this in cane toads: under dehydrating conditions, wild-caught individuals from Australia exhibit better locomotor performance than do conspecifics from Hawai’i and the native range (G. Kosmala, Christian, Brown, & Shine, 2017), and within Australia, toads from semi-arid areas (i.e. NT) exhibit better locomotor performance than do conspecifics from wetter areas (i.e. coastal QLD) (G. Kosmala et al., 2017; Tingley, Greenlees, & Shine, 2012). This trait is heritable; captive-bred toads with parental origins from hotter areas (northwestern Australia) outperform those with parental origins from cooler areas (northeastern Australia) at high (but not low) temperatures in a common-garden setting (G. K. Kosmala, Brown, Christian, et al., 2018). Coupled with the previously shown heritable phenotypic patterns, our results suggest that intermediate and frontal toads in Australia may be evolving enhanced thermal tolerance, thereby facilitating their continued westward range expansion (Szucs et al., 2017). However, manipulative experiments are required to directly test this hypothesis.

Although our data suggest that invasive toads may be responding to localized natural selection, our data also support the presence of other evolutionary forces. We found evidence of isolation by distance (IBD) in Australia using the full SNP dataset, which is likely to be caused by genetic drift. However, loci under strong selection are less likely to show evidence of IBD. Unsurprisingly, when we plot genetic vs. geographic distance in Australia, using loci putatively associated with environmental selection, we see a clear discontinuity in these data (Fig 4C). Other evolutionary forces may be more difficult to detect in these data. Spatial sorting is the separation of individuals within a range-expanding population along phases of their expansion based on their dispersive capabilities, and was first characterized in Australian cane toads (Shine et al., 2011). Spatial sorting is expected to result in clinal variation in dispersal-related allele frequencies along the range, so we might expect that the relationship between genetic and geographic distance using loci underlying dispersal-related characteristics would show a similar pattern to that of genetic drift. In this study, when we examine this relationship only using loci identified as under selection in this introduction, but not associated with environmental selection, there is a gradual decay of the relationship between genetic and geographic distance (Fig 4D). Although we acknowledge that these loci may be influenced by drift to some degree, it seems plausible that because they were identified as FST outliers, they are likely to be influenced by other evolutionary forces, including spatial sorting.

Our population structure analyses using the RADSeq dataset (Trumbo et al., 2016) were similar to those from our RNA-Seq dataset; both showed two genetic clusters within Australia. However, unlike the RNA-Seq results (which show a divide occurring in inland QLD, where temperature increases and rainfall decreases), the RADSeq results cluster all QLD toads in the first group, and NT/WA toads in the second group. In the RADSeq dataset, the spatial location of the transition between the two groups is difficult to pinpoint due to the absence of intermediate sampling sites. The difference in group divisions between the two datasets may be because selection from environmental variables causes differentiation between toads from coastal and inland QLD (as seen in the RNA-Seq data), whereas demographic processes such as gene flow may lessen differentiation between them (as seen in the RADSeq data). Alternatively, toads from coastal and inland QLD may cluster together in the RADSeq data because the sampling sites in that study were south of those in our study, and the environmental conditions of coastal and inland sites are more similar in that area (Hijmans et al., 2005). Estimates of diversity from the RADSeq dataset were slightly higher than those from the RNA-Seq dataset; this may be because adaptive variation is expected to be lower than neutral variation due to evolutionary constraints (C. A. Andrews, 2010).

In conclusion, it appears that serial introductions have led to a reduction in genetic diversity of invasive cane toads. Nonetheless, cane toads do indeed face novel adaptive challenges in their introduced ranges, and are able to respond genetically to natural selection, particularly in the harsh environmental conditions of northwestern Australia. The ability of toads to adapt to these conditions may reflect the maintenance of diversity at ecologically relevant loci, or it may reflect sufficiently high mutation rates to bolster adaptive potential; thus, it remains unclear whether the toad invasion represents a true genetic paradox. We have found evidence of natural selection, genetic drift, and spatial sorting in the Australian invasion, but the impacts of these forces require further investigation to tease apart. Manipulative experiments investigating the candidates that we have identified may be especially informative. Additionally, genetic changes may not be the only mechanisms by which adaptation occurs; studies on plasticity and epigenetic changes may also be useful for uncovering the mysteries of rapid evolution.

## Acknowledgements

This work was supported by the Australian Research Council (FL120100074, DE150101393) and the Equity Trustees Charitable Foundation (Holsworth Wildlife Research Endowment). We thank Olivier Francois and Natalie Hofmeister for their input during discussions about our analyses. We thank Cam Hudson, Joachim Ehlenz, Serena Lam, and Chris Jolly for their assistance with sample collection. We thank Andrea West for her assistance during RNA extractions. All procedures involving live animals were approved by the University of Sydney Animal Care and Ethics Committee.

