## Supplementary material for "Bottleneck revisited: increased adaptive variation despite reduced overall genetic diversity in a rapidly adapting invader": Fig S1

Invaders weather the weather: rapid adaptation to a novel environment despite evidence of genetic drift and spatial sorting

**Supplemental Information**

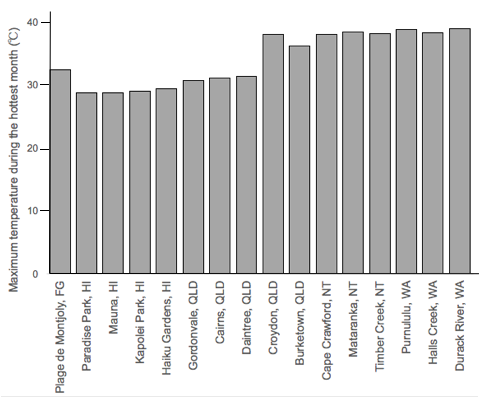

Fig S1. Maximum temperature during the warmest month in each location across collection sites for invasive cane toads in French Guiana, Hawaii, and Australia.

Table S1.

(A) Collection sites for an RNA-Seq experiment on spleen tissue from invasive cane toads from French Guiana, Hawaii, and Australia (N=8 native range, N=10 source, N=10 core, N=8 intermediate, N=10 front).

| Site | Phase | Coordinates |
| --- | --- | --- |
| Plage de Montjoly, French Guiana | Native Range | 4.91327, -52.2599 |
| Kapolei Regional Park, Oahu | Source | 21.3356, -158.0779 |
| Haiku Gardens, Oahu | Source | 21.4151, -157.8157 |
| Gordonvale, QLD | Core | -17.0832, 145.7961 |
| Daintree, QLD | Core | -16.25, 145.3167 |
| Cape Crawford, NT | Intermediate | -16.6667, 135.8 |
| Timber Creek, NT | Intermediate | -15.6432, 130.4666 |
| Halls Creek, WA | Front | -18.2247, 127.6729 |
| Durack River, WA | Front | -15.9068, 128.184 |

(B) Collection sites for an RNA-Seq experiment on brain tissue from invasive cane toads from Hawaii and Australia (N=18 for each phase of the invasion).

| Site | Phase | Coordinates |
| --- | --- | --- |
| Paradise Park, Big Island | Source | 19.5724, -154.9577 |
| Mauna Lani Point, Big Island | Source | 19.9385, -155.8754 |
| Kapolei Regional Park, Oahu | Source | 21.3356, -158.0779 |
| Haiku Gardens, Oahu | Source | 21.4151, -157.8157 |
| Gordonvale, QLD | Core | -17.0832, 145.7961 |
| Cairns, QLD | Core | -16.9186, 145.7781 |
| Daintree, QLD | Core | -16.25, 145.3167 |
| Croydon, QLD | Intermediate | -18.206, 142.24 |
| Burketown, QLD | Intermediate | -17.8522, 139.633 |
| Cape Crawford, NT | Intermediate | -16.6667, 135.8 |
| Mataranka, NT | Intermediate | -14.9234, 133.066 |
| Timber Creek, NT | Intermediate | -15.6432, 130.4666 |
| Purnululu, WA | Front | -17.4484, 128.5467 |
| Halls Creek, WA | Front | -18.2247, 127.6729 |
| Durack River, WA | Front | -15.9068, 128.184 |

Table S2.

(A) SNPs with outlier F_ST_ values and significant associations with maximum temperature during the warmest month of the year (per collection site) from RNA-Seq data from spleens in cane toads from French Guiana, Hawaii, and Australia.

| gene | base position | protein |
| --- | --- | --- |
| *CDK12* | 3267 | Cyclin-dependent kinase 12 |
| *TRM1L* | 966 | TRMT1-like protein |
| *THOC1* | 2284 | THO complex subunit 1 |
| *TRM1L* | 2088 | TRMT1-like protein |
| *HSPA4* | 2006 | Heat shock 70 kDa protein 4 |
| *TRM1L* | 951 | TRMT1-like protein |
| *HSPA4* | 1043 | Heat shock 70 kDa protein 4 |
| *SMG1* | 9837 | Serine/threonine-protein kinase SMG1 |
| *NFIP1* | 2254 | NEDD4 family-interacting protein 1 |
| *NFIP1* | 1188 | NEDD4 family-interacting protein 1 |
| *DREB* | 574 | Drebrin |
| *CYLD* | 871 | Ubiquitin carboxyl-terminal hydrolase CYLD |
| *TRM1L* | 2100 | TRMT1-like protein |
| *TYSY* | 758 | Thymidylate synthase |
| *PIGT* | 1937 | GPI transamidase component PIG-T |
| *TYSY* | 677 | Thymidylate synthase |
| *FUBP3* | 1638 | Far upstream element-binding protein 3 |
| *NFIP1* | 2629 | NEDD4 family-interacting protein 1 |
| *NFIP1* | 1725 | NEDD4 family-interacting protein 1 |
| *TOB1* | 882 | Protein Tob1 |
| *UBF1B* | 2159 | Nucleolar transcription factor 1-B |
| *NMRL1* | 129 | NmrA-like family domain-containing protein 1 |
| *YES* | 2756 | Tyrosine-protein kinase Yes |
| *TCPZ* | 100 | T-complex protein 1 subunit zeta |
| *NOMO3* | 743 | Nodal modulator 3 |
| *TYSY* | 800 | Thymidylate synthase |
| *VATO* | 538 | V-type proton ATPase 21 kDa proteolipid subunit |
| *PYR1* | 6103 | CAD protein |
| *MANEA* | 1162 | Glycoprotein endo-alpha-1,2-mannosidase |
| *ZBT7B* | 1709 | Zinc finger and BTB domain-containing protein 7B |
| *AL9A1* | 588 | 4-trimethylaminobutyraldehyde dehydrogenase |
| *AL9A1* | 582 | 4-trimethylaminobutyraldehyde dehydrogenase |
| *SON* | 12805 | Protein SON |
| *SBDS* | 2613 | Ribosome maturation protein SBDS |
| *VATO* | 368 | V-type proton ATPase 21 kDa proteolipid subunit |
| *SON* | 9429 | Protein SON |
| *PYR1* | 6973 | CAD protein |
| *PTN6* | 2024 | Tyrosine-protein phosphatase non-receptor type 6 |
| *WLSB* | 292 | Protein wntless homolog B |
| *PTN6* | 2362 | Tyrosine-protein phosphatase non-receptor type 6 |
| *PYR1* | 5474 | CAD protein |
| *TCRG1* | 957 | Transcription elongation regulator 1 |
| *SON* | 16145 | Protein SON |
| *AL9A1* | 578 | 4-trimethylaminobutyraldehyde dehydrogenase |
| *DHX8* | 1534 | ATP-dependent RNA helicase DHX8 |
| *STRN* | 3715 | Striatin |
| *RBM34* | 276 | RNA-binding protein 34 |
| *WLSB* | 357 | Protein wntless homolog B |
| *PYR1* | 1993 | CAD protein |
| *AT11C* | 6666 | Phospholipid-transporting ATPase IG |
| *PAXB1* | 2726 | PAX3- and PAX7-binding protein 1 |
| *SAP* | 1499 | Prosaposin |
| *SON* | 12673 | Protein SON |
| *2AAA* | 964 | Serine/threonine-protein phosphatase 2A 65 kDa regulatory subunit A alpha isoform |
| *PRLD1* | 330 | PRELI domain-containing protein 1, mitochondrial |
| *PRLD1* | 70 | PRELI domain-containing protein 1, mitochondrial |
| *.* | 733 | . |
| *DDX41* | 2017 | Probable ATP-dependent RNA helicase DDX41 |
| *ODO1* | 3515 | 2-oxoglutarate dehydrogenase, mitochondrial |
| *LEMD2* | 928 | LEM domain-containing protein 2 |
| *TTC31* | 1157 | Tetratricopeptide repeat protein 31 |
| *SYNRG* | 2895 | Synergin gamma |
| *LEMD2* | 2168 | LEM domain-containing protein 2 |
| *NBR1* | 3313 | Next to BRCA1 gene 1 protein |

(B) SNPs with outlier F_ST_ values and significant associations with maximum temperature during the warmest month of the year (per collection site) from RNA-Seq data from brains in invasive cane toads from Hawaii and Australia.

| gene | base position | protein |
| --- | --- | --- |
| *RASA1* | 3118 | Ras GTPase-activating protein 1 |
| *SYLC* | 3425 | Leucine--tRNA ligase, cytoplasmic |
| *NFIP1* | 1452 | NEDD4 family-interacting protein 1 |
| *NFIP1* | 2703 | NEDD4 family-interacting protein 1 |
| *NFIP1* | 3135 | NEDD4 family-interacting protein 1 |
| *NFIP1* | 360 | NEDD4 family-interacting protein 1 |
| *NFIP1* | 915 | NEDD4 family-interacting protein 1 |
| *NFIP1* | 1188 | NEDD4 family-interacting protein 1 |
| *NFIP1* | 1725 | NEDD4 family-interacting protein 1 |
| *NFIP1* | 1942 | NEDD4 family-interacting protein 1 |
| *NFIP1* | 2214 | NEDD4 family-interacting protein 1 |
| *NFIP1* | 2254 | NEDD4 family-interacting protein 1 |
| *NFIP1* | 2629 | NEDD4 family-interacting protein 1 |
| *SMG9* | 1270 | Protein SMG9 |
| *SMG9* | 1285 | Protein SMG9 |
| *HSPA4* | 2688 | Heat shock 70 kDa protein 4 |
| *DREB* | 574 | Drebrin |
| *RNF14* | 1152 | E3 ubiquitin-protein ligase RNF14 |
| *PRELID1* | 70 | PRELI domain-containing protein 1, mitochondrial |
| *PRELID1* | 330 | PRELI domain-containing protein 1, mitochondrial |
| *RNF14* | 1757 | E3 ubiquitin-protein ligase RNF14 |
| *AT2B4* | 8002 | Plasma membrane calcium-transporting ATPase 4 |
| *AT2B4* | 8123 | Plasma membrane calcium-transporting ATPase 4 |
| *QCR2* | 1954 | Cytochrome b-c1 complex subunit 2, mitochondrial |
| *DDX41* | 2017 | Probable ATP-dependent RNA helicase DDX41 |
| *-* | 596 | - |
| *QCR2* | 1987 | Cytochrome b-c1 complex subunit 2, mitochondrial |
| *ARNT2* | 1008 | Aryl hydrocarbon receptor nuclear translocator 2 |
| *BDP1* | 383 | Transcription factor TFIIIB component B'' homolog |
| *QCR2* | 1952 | Cytochrome b-c1 complex subunit 2, mitochondrial |
| *SERF1* | 378 | Small EDRK-rich factor 1 |
| *QCR2* | 2010 | Cytochrome b-c1 complex subunit 2, mitochondrial |
| *QCR2* | 2038 | Cytochrome b-c1 complex subunit 2, mitochondrial |
| *RUVB1* | 2921 | RuvB-like 1 |
| *MO4L1* | 2128 | Mortality factor 4-like protein 1 |
| *AB17C* | 443 | Protein ABHD17C |
| *PIGV* | 2619 | GPI mannosyltransferase 2 |
| *AN32E* | 491 | Acidic leucine-rich nuclear phosphoprotein 32 family member E |

Table S3.

(A) SNPs with outlier F_ST_ values and significant associations with rainfall during the driest quarter of the year (per collection site) from RNA-Seq data from spleen in cane toads from French Guiana, Hawaii, and Australia.

| gene | base position | protein |
| --- | --- | --- |
| *PTPRC* | 6803 | Receptor-type tyrosine-protein phosphatase C |
| *TPRN* | 2983 | Taperin |
| *TPRN* | 3800 | Taperin |
| *PCNT* | 12271 | Pericentrin |
| *TCPD* | 2105 | T-complex protein 1 subunit delta |
| *ANR10* | 1164 | Ankyrin repeat domain-containing protein 10 |
| *B3GN2* | 1554 | N-acetyllactosaminide beta-1,3-N-acetylglucosaminyltransferase 2 |
| *SMG1* | 9837 | Serine/threonine-protein kinase SMG1 |
| *GANP* | 5116 | Germinal-center associated nuclear protein |
| *GANP* | 7035 | Germinal-center associated nuclear protein |
| *PCNT* | 12168 | Pericentrin |
| *GANP* | 5852 | Germinal-center associated nuclear protein |
| *SFRP3* | 1044 | Secreted frizzled-related protein 3 |
| *MYPOP* | 1291 | Myb-related transcription factor, partner of profilin |
| *PCNT* | 13249 | Pericentrin |
| *SMUF2* | 3385 | E3 ubiquitin-protein ligase SMURF2 |
| *PCNT* | 12106 | Pericentrin |
| *GANP* | 6952 | Germinal-center associated nuclear protein |
| *LHFP* | 2029 | Lipoma HMGIC fusion partner |
| *2AAA* | 2017 | Serine/threonine-protein phosphatase 2A 65 kDa regulatory subunit A alpha isoform |
| *TOB1* | 882 | Protein Tob1 |
| *BECN1* | 1151 | Beclin-1 |
| *BTBDA* | 1531 | BTB/POZ domain-containing protein 10 |
| *.* | 639 | . |
| *NMRL1* | 129 | NmrA-like family domain-containing protein 1 |
| *SREK1* | 1694 | Splicing regulatory glutamine/lysine-rich protein 1 |
| *TCPQ* | 971 | T-complex protein 1 subunit theta |
| *TCPQ* | 1088 | T-complex protein 1 subunit theta |
| *VATO* | 538 | V-type proton ATPase 21 kDa proteolipid subunit |
| *PYR1* | 6103 | CAD protein |
| *AL9A1* | 588 | 4-trimethylaminobutyraldehyde dehydrogenase |
| *AL9A1* | 582 | 4-trimethylaminobutyraldehyde dehydrogenase |
| *SON* | 12805 | Protein SON |
| *VATO* | 368 | V-type proton ATPase 21 kDa proteolipid subunit |
| *SON* | 9429 | Protein SON |
| *PYR1* | 6973 | CAD protein |
| *PYR1* | 5474 | CAD protein |
| *SON* | 16145 | Protein SON |
| *AL9A1* | 578 | 4-trimethylaminobutyraldehyde dehydrogenase |
| *NUBP2* | 834 | Cytosolic Fe-S cluster assembly factor nubp2 |
| *NUBP2* | 970 | Cytosolic Fe-S cluster assembly factor nubp2 |
| *PYR1* | 1993 | CAD protein |
| *NUBP2* | 959 | Cytosolic Fe-S cluster assembly factor nubp2 |
| *SON* | 12673 | Protein SON |
| *MRP3* | 4668 | Canalicular multispecific organic anion transporter 2 |
| *SYNRG* | 2895 | Synergin gamma |
| *LEMD2* | 2168 | LEM domain-containing protein 2 |

(B) SNPs with outlier F_ST_ values and significant associations with rainfall during the driest quarter of the year (per collection site) from RNA-Seq data from brains in invasive cane toads from Hawaii and Australia.

| gene | base position | protein |
| --- | --- | --- |
| *MAGED2* | 1606 | Melanoma-associated antigen D2 |
| *MAGED2* | 1608 | Melanoma-associated antigen D2 |
| *MAGED2* | 1613 | Melanoma-associated antigen D2 |
| *MAGED2* | 1057 | Melanoma-associated antigen D2 |
| *MAGED2* | 1065 | Melanoma-associated antigen D2 |
| *MAGED2* | 1068 | Melanoma-associated antigen D2 |
| *MAGED2* | 1071 | Melanoma-associated antigen D2 |
| *MAGED2* | 1074 | Melanoma-associated antigen D2 |
| *MAGED2* | 1097 | Melanoma-associated antigen D2 |
| *MAGED2* | 1492 | Melanoma-associated antigen D2 |
| *MAGED2* | 1516 | Melanoma-associated antigen D2 |
| *MAGED2* | 1522 | Melanoma-associated antigen D2 |
| *MAGED2* | 1577 | Melanoma-associated antigen D2 |
| *MAGED2* | 1581 | Melanoma-associated antigen D2 |
| *MAGED2* | 1033 | Melanoma-associated antigen D2 |
| *RBM34* | 1282 | RNA-binding protein 34 |
| *EIF3A* | 834 | Eukaryotic translation initiation factor 3 subunit A |
| *MAGED2* | 982 | Melanoma-associated antigen D2 |
| *BAP31* | 261 | B-cell receptor-associated protein 31 |
| *TOB1* | 882 | Protein Tob1 |
| *TOB1* | 1299 | Protein Tob1 |
| *MAGED2* | 2045 | Melanoma-associated antigen D2 |
| *MAGED2* | 2046 | Melanoma-associated antigen D2 |
| *TF3C5* | 1136 | General transcription factor 3C polypeptide 5 |
| *MAGED2* | 1998 | Melanoma-associated antigen D2 |
| *STXBP1* | 147 | Syntaxin-binding protein 1 |
| *STXBP1* | 1672 | Syntaxin-binding protein 1 |
| *STXBP1* | 2752 | Syntaxin-binding protein 1 |
| *STXBP1* | 3135 | Syntaxin-binding protein 1 |
| *SMC1A* | 2486 | Structural maintenance of chromosomes protein 1A |
| *SMC1A* | 2522 | Structural maintenance of chromosomes protein 1A |
| *SMC1A* | 2816 | Structural maintenance of chromosomes protein 1A |
| *GNDS* | 3242 | Ral guanine nucleotide dissociation stimulator |

Table S4.

(A) Pairwise F_ST_ by invasion phase based on SNPs in RNA-Seq data from brains in invasive cane toads from Hawaii and Australia. Source = Hawaii; Core = QLD, Australia; Intermediate = NT, Australia; Front = WA, Australia. N=18 for each phase.

|  | Source | Core | Intermediate | Front |
| --- | --- | --- | --- | --- |
| Source |  |  |  |  |
| Core | 0.04 |  |  |  |
| Intermediate | 0.08 | 0.06 |  |  |
| Front | 0.11 | 0.08 | 0.01 |  |

(B) Pairwise F_ST_ by collection site based on SNPs in RNA-Seq data from brains in invasive cane toads from Hawaii and Australia

|  | Paradise Park | Mauna Kea | Kapolei Regional Park | Haiku Gardens | Gordonvale | Cairns | Daintree | Croydon | Burketown | Cape Crawford | Mataranka | Timber Creek | Purnululu | Halls Creek | Durack River |
| --- | --- | --- | --- | --- | --- | --- | --- | --- | --- | --- | --- | --- | --- | --- | --- |
| Paradise Park |  |  |  |  |  |  |  |  |  |  |  |  |  |  |  |
| Mauna Kea | 0.09 |  |  |  |  |  |  |  |  |  |  |  |  |  |  |
| Kapolei Regional Park | 0.06 | 0.10 |  |  |  |  |  |  |  |  |  |  |  |  |  |
| Haiku Gardens | 0.06 | 0.10 | 0.07 |  |  |  |  |  |  |  |  |  |  |  |  |
| Gordonvale | 0.07 | 0.09 | 0.08 | 0.06 |  |  |  |  |  |  |  |  |  |  |  |
| Cairns | 0.05 | 0.08 | 0.07 | 0.06 | 0.01 |  |  |  |  |  |  |  |  |  |  |
| Daintree | 0.07 | 0.11 | 0.09 | 0.08 | 0.04 | 0.03 |  |  |  |  |  |  |  |  |  |
| Croydon | 0.09 | 0.10 | 0.10 | 0.09 | 0.05 | 0.05 | 0.08 |  |  |  |  |  |  |  |  |
| Burketown | 0.10 | 0.12 | 0.12 | 0.10 | 0.05 | 0.06 | 0.08 | 0.03 |  |  |  |  |  |  |  |
| Cape Crawford | 0.11 | 0.12 | 0.12 | 0.11 | 0.06 | 0.07 | 0.09 | 0.03 | 0.01 |  |  |  |  |  |  |
| Mataranka | 0.11 | 0.13 | 0.13 | 0.11 | 0.06 | 0.07 | 0.08 | 0.03 | 0.01 | -0.01 |  |  |  |  |  |
| Timber Creek | 0.12 | 0.16 | 0.14 | 0.12 | 0.08 | 0.09 | 0.10 | 0.06 | 0.03 | 0.00 | -0.00 |  |  |  |  |
| Purnululu | 0.13 | 0.14 | 0.14 | 0.13 | 0.08 | 0.09 | 0.10 | 0.05 | 0.03 | 0.01 | 0.01 | 0.00 |  |  |  |
| Halls Creek | 0.13 | 0.16 | 0.15 | 0.14 | 0.09 | 0.10 | 0.10 | 0.06 | 0.03 | 0.01 | 0.01 | 0.00 | -0.00 |  |  |
| Durack River | 0.12 | 0.15 | 0.14 | 0.12 | 0.07 | 0.08 | 0.10 | 0.05 | 0.02 | -0.00 | -0.00 | 0.00 | 0.00 | 0.00 |  |

Table S5. Expected heterozygosity (He), allelic richness (AR), and Shannon’s Information Index (SI) estimated using SNPs from spleen RNA-Seq data on native and invasive populations of cane toads (*Rhinella marina*) at collection sites across French Guiana, Hawai’i, and Australia.

| Collection site | All loci  (He, AR, SI) | Outlier F_ST_ loci  (He, AR, SI) | Environmentally associated loci  (He, AR, SI) |
| --- | --- | --- | --- |
| French Guiana (native, N=8) | 0.27, 1.66, 0.40 | 0.03, 1.25, 0.06 | 0.17, 1.53, 0.25 |
| Kapolei Park (source, N=5) | 0.23, 1.60, 0.34 | 0.07, 1.20, 0.10 | 0.28, 1.77, 0.41 |
| Haiku Gardens (source, N=5) | 0.23, 1.60, 0.34 | 0.07, 1.22, 0.11 | 0.28, 1.75, 0.41 |
| Gordonvale (core, N=5) | 0.24, 1.63, 0.36 | 0.07, 1.21, 0.10 | 0.30, 1.84, 0.45 |
| Daintree (core, N=5) | 0.24, 1.62, 0.35 | 0.08, 1.24, 0.12 | 0.29, 1.79, 0.44 |
| Cape Crawford (intermediate, N=4) | 0.22, 1.59, 0.32 | 0.02, 1.06, 0.03 | 0.22, 1.59, 0.32 |
| Timber Creek (intermediate, N=4) | 0.21, 1.58, 0.32 | 0.02, 1.05, 0.03 | 0.22, 1.60, 0.32 |
| Halls Creek (front, N=5) | 0.21, 1.57, 0.32 | 0.02, 1.05, 0.03 | 0.19, 1.58, 0.30 |
| Durack River (front, N=5) | 0.22, 1.58, 0.32 | 0.03, 1.11, 0.05 | 0.20, 1.60, 0.31 |

Table S6. Expected heterozygosity (He), allelic richness (AR), and Shannon’s Information Index (SI) estimated using SNPs from brain RNA-Seq data on native and invasive populations of cane toads (*Rhinella marina*) at collection sites across Hawai’i and Australia.

| Collection site | All loci  (He, AR, SI) | Outlier F_ST_ loci  (He, AR, SI) | Environmentally associated loci  (He, AR, SI) |
| --- | --- | --- | --- |
| Paradise Park (source, N=4) | 0.28, 1.57, 0.41 | 0.31, 1.76, 0.45 | 0.30, 1.84, 0.45 |
| Mauna Lani (source, N=4) | 0.27, 1.56, 0.40 | 0.24, 1.61, 0.35 | 0.29, 1.73, 0.42 |
| Kapolei Park (source, N=5) | 0.29, 1.57, 0.43 | 0.29, 1.83, 0.43 | 0.25, 1.73, 0.38 |
| Haiku Gardens (source, N=5) | 0.30, 1.59, 0.44 | 0.31, 1.75, 0.45 | 0.31, 1.83, 0.46 |
| Gordonvale (core, N=4) | 0.30, 1.61, 0.44 | 0.36, 1.82, 0.51 | 0.22, 1.62, 0.33 |
| Cairns (core, N=10) | 0.32, 1.60, 0.48 | 0.38, 1.95, 0.55 | 0.24, 1.84, 0.38 |
| Daintree (core, N=4) | 0.29, 1.60, 0.43 | 0.32, 1.90, 0.48 | 0.18, 1.57, 0.28 |
| Croydon (intermediate, N=2) | 0.26, 1.60, 0.37 | 0.18, 1.45, 0.26 | 0.10, 1.24, 0.14 |
| Burketown (intermediate, N=4) | 0.28, 1.57, 0.41 | 0.18, 1.44, 0.26 | 0.09, 1.31, 0.14 |
| Cape Crawford (intermediate, N=4) | 0.28, 1.58, 0.42 | 0.15, 1.42, 0.23 | 0.08, 1.32, 0.13 |
| Mataranka (intermediate, N=4) | 0.28, 1.57, 0.41 | 0.16, 1.46, 0.24 | 0.07, 1.25, 0.11 |
| Timber Creek (intermediate, N=4) | 0.27, 1.55, 0.40 | 0.05, 1.18, 0.09 | 0.06, 1.22, 0.10 |
| Purnululu (front, N=10) | 0.29, 1.55, 0.44 | 0.20, 1.48, 0.12 | 0.07, 1.37, 0.11 |
| Halls Creek (front, N=4) | 0.27, 1.56, 0.40 | 0.09, 1.25, 0.14 | 0.06, 1.21, 0.10 |
| Durack River (front, N=4) | 0.27, 1.56, 0.41 | 0.13, 1.37, 0.19 | 0.06, 1.24, 0.10 |

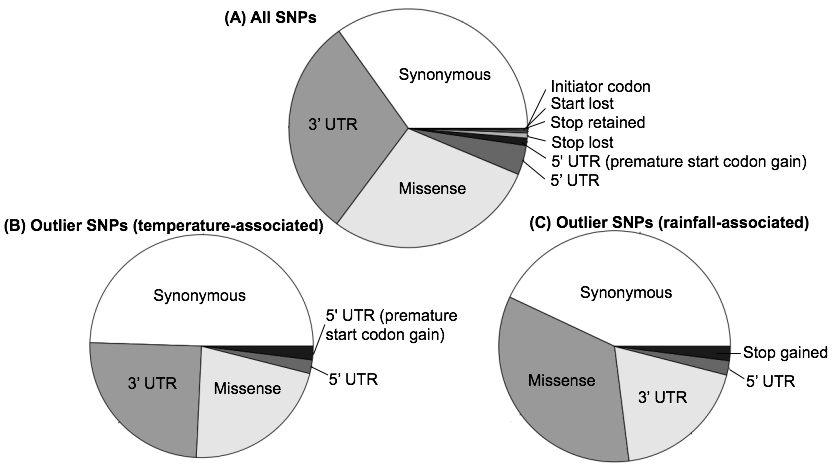

Fig S2. Types of SNPs and in RNA-Seq data from spleens in cane toads across sites in French Guiana, Hawai’i, and Australia. (A) represents the full dataset. (B) represents only F_ST_ outliers with an association with maximum temperature during the hottest month. (C) represents only F_ST_ outliers with an association with rainfall during the driest quarter.

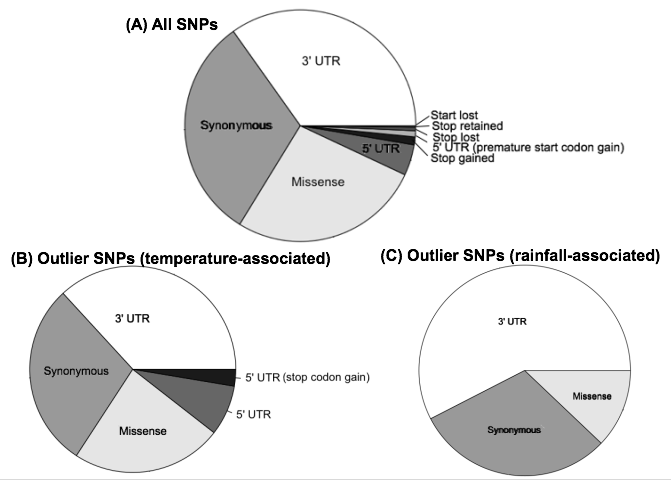

FigS3. Types of SNPs and in RNA-Seq data from brains in invasive cane toads across sites in Hawai’i and Australia. (A) represents the full dataset. (B) represents only F_ST_ outliers with an association with maximum temperature during the hottest month. (C) represents only F_ST_ outliers with an association with rainfall during the driest quarter.
